# *IN SILICO* DRUG DISCOVERY FOR NDM-1 METALLO β LACTAMASE INHIBITORS for *KLEBSIELLA PUEUMONIAE*

**DOI:** 10.1101/2025.08.16.670670

**Authors:** S Raghavendra, P Rashmi, E Prashanth, A Yashwanth Bharadwaj, E Ranjith, S Varun, VG Shanmuga Priya

## Abstract

In every region of the world, antibiotic resistance is increasing to dangerously high levels. Our ability to cure widespread infectious diseases is being threatened by the emergence of new resistance mechanisms*. Klebsiella pneumoniae* is one of the most prevalent nosocomial Gram-negative bacteria in the world. NDM-1 is a brand-new class of metallo-β-lactamase (MBL) that makes the bacteria almost total resistant to all β-lactam antibiotics, including penicillins, cephalosporins and carbapenems. Dangerous infections could develop if NDM-1 switches to a bacterium that is already resistant to antibiotics. It could be untreatable and spread quickly among humans. Using the available insilico tools, in the present research work, an attempt has been made to develop inhibitors for NDM-1 β Lactamase of *K. pneumoniae*. In the present study, 8 standard ligands were identified and docked against the NDM-1 protein using PyRx. Among these standards, Sulfonamide was selected as the best compound and virtual screening of a large number of sulfonamide moieties was conducted with the identified active site in NDM-1 protein using PyRx. On analyzing the obtained results, about 60 molecules were selected as best hits for Docking studies and ADMET studies. On analyzing the ADMET properties and binding energies of the top hits, (3Z)-N-hydroxypenta-1,3-diene-2-sulfonamide (Pubchem CID 118156306) and N-hydroxyfuran-2-sulfonamide (Pubchem CID 46175386). Molecular dynamics simulations were carried out on the complxes formed by these 2 compounds with target protein and the results showed the complexes were stable which validated the earlier findings. Based on the findings of the study, it was concluded that (3Z)-N-hydroxypenta-1,3-diene-2-sulfonamide (Pubchem CID 118156306) and N-hydroxyfuran-2-sulfonamide (Pubchem CID 46175386) had the potential to be used as lead candidates against infections caused by *K. pneumoniae* producing NDM-1 strains.

## 1. INTRODUCTION

Our ability to cure widespread infectious diseases is being threatened by the emergence and global dissemination of new resistance mechanisms in microorganisms. One of the largest risks to food security, and global health is antibiotic resistance. Modern medicine’s advancements are under jeopardy due to antibiotic resistance. Without efficient antibiotics, prevention and treatment of infections during organ transplants, chemotherapy, and procedures like caesarean sections become significantly more risky^1^. In 2009, a strain of *Klebsiella pneumoniae* bacteria with broad-spectrum antibiotic resistance was discovered and isolated in a Swedish patient, formerly hospitalized in India. The antibiotic resistance determinant was recognized as a novel Metallo-β-lactamase and designated as NDM-1(*bla*_NDM-1_ gene), which is short form of New Delhi metallo-β-lactamase^2^. NDM-1 is a brand-new class of metallo-β-lactamase (MBL) that makes the bacteria almost total resistant to all β-lactam antibiotics, including penicillins, cephalosporins and carbapenems^3^. NDM-1 is seen expressed in organisms causing pneumonia, urinary tract infections, and intra-abdominal infections. It ranks among the top three causal agents in most situations, making it a significant contributor to newborn sepsis. The World Health Organization (WHO) is concerned about NDM-1’s potential to see “the doomsday scenario of a world without antibiotics.”

**Fig 1.**
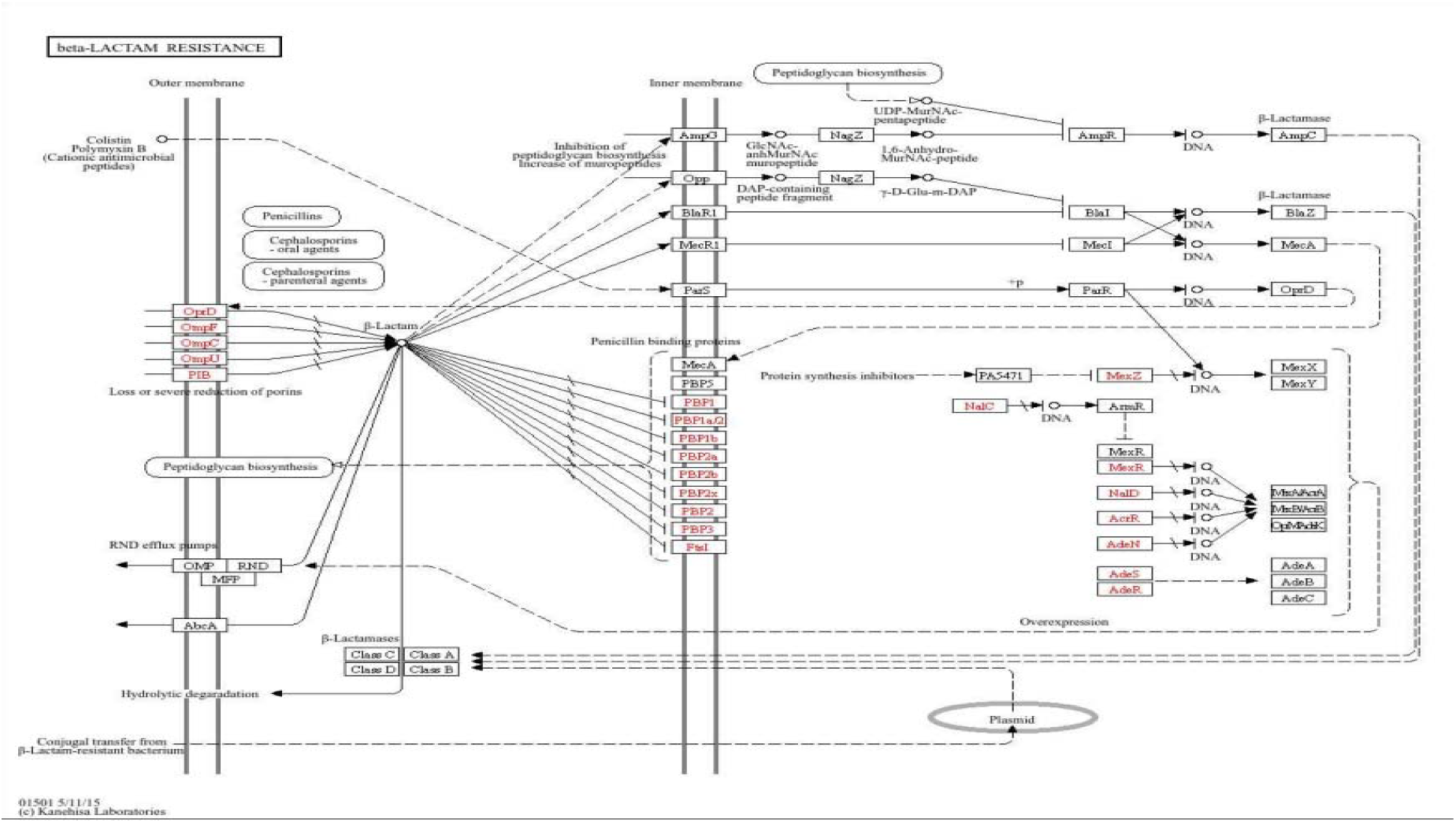
Pathway of NDM causing Antibiotic Resistance

The *bla*_NDM-1_ gene produces Carbapenem’s-β-lactamase that causes hydrolytic degradation of β-lactam ring by using 2 zinc ions found in active site of NDM-1 which form Zn-H coordination bond on hydrolysis which inhibits production of penicillin binding proteins which are essential for antibiotic action thus making the antibiotics ineffective.^4^

NDM-1 can spread to other organisms by Horizontal gene transfer (HGT) where *bla*_NDM-_1 gene is transferred to other organisms through plasmids making the bacteria that receives the gene also resistant to antibiotics through this mechanism. This was discovered in United Kingdom when NDM-1 gene was found *Escherichia Coli.* This makes NDM-1 more dangerous as it can spread across organisms^5^. If NDM-1 spreads to other microbes, additional illnesses will develop and cause a global health emergency.

So far, Patients with NDM-1 related illnesses have received case-by-case treatment using a mix of drugs like colistin, succinic and oxalic acid. Undoubtedly, finding new NDM-1 targeting medication candidates is extremely important for battling disorders caused by NDM-1^6^.

By the studies that were already performed it was noticed that mostly-β-lactam antibiotics are only being used for treatment of infections caused by organisms expressing NDM-1. It was surprising to find that sulfonamide has the properties to inhibit NDM-1 strains which is further explored in this study to formulate a proper drug to treat the infections caused by NDM-1^7–8^.

## 2. METHODOLOGY

### 2.1 RETRIEVAL OF TARGET PROTEIN AND SMALL MOLECULES

Protein NDM-1 with a ligand (Crystal Structure of NDM-1 D199N with Compound 16) was obtained from PDB Database (PDB id: 6NY7) (uniprot id-<u>C7C422</u>). **(5,7-dibromo-2-oxo-1,2-dihydroquinolin-4-yl) methyl phosphonic acid** was the native ligand found in the protein 6NY7^9^. By studying interactions, an active site was identified as target in the protein.

Pubchem Database was used to retrieve small molecules such as Penicillin, Benzyl Penicillin, Mercaptopurine, Ampicillin, Sulfonamide, cyclobutanone, Triazole and Quercitin. As this study is focused on sulfonamide, Lipinski and verber’s rule were used to filter and retrieve about 1400 compounds from all the sulfonamides available in pubchem database and was retrieved as single file (SDF Format) which was converted to PDB format using Openbabel for further studies.^10^ (file attached in supplement)

### 2.2 DOCKING STUDIES

The obtained NDM-1 protein (6ny7) structure was subjected to purification to make it suitable for docking using Autodock tools 1.5.7, where the native ligand, water and other foreign entities in the protein structure were removed followed by adding kollman and gasteiger charges. The native ligand (Pubchem CID:139033900) was docked again to the identified active site using Autodock Vina and the obtained interactions were found to be same as shown in PDB database. Thus, our docking model was validated. (Results shown in supplement table). Docking studies of Standard ligands retrieved from Pubchem with active site of NDM-1 protein was performed using Autodock Vina^11–13^.

**Table 1:**
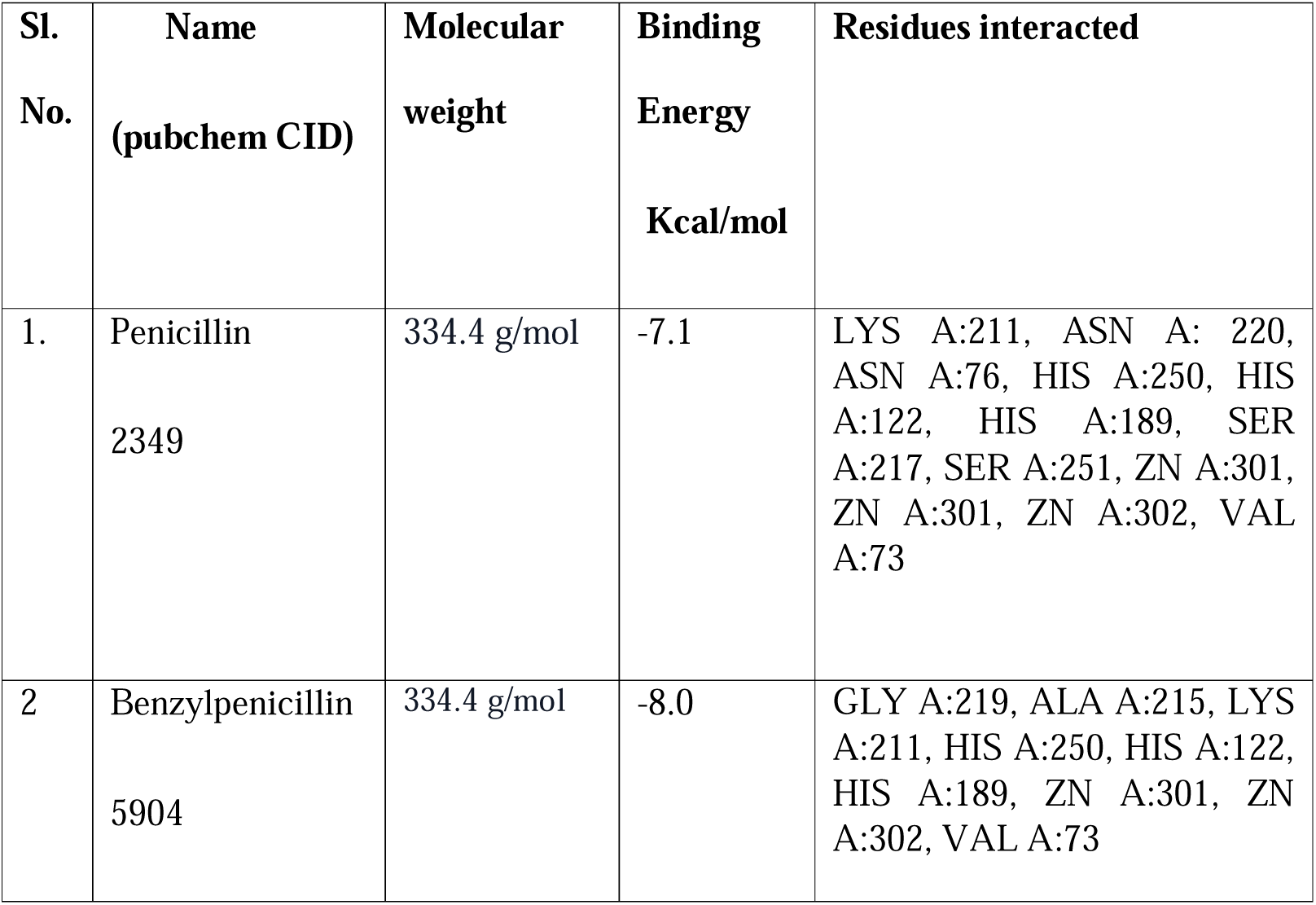
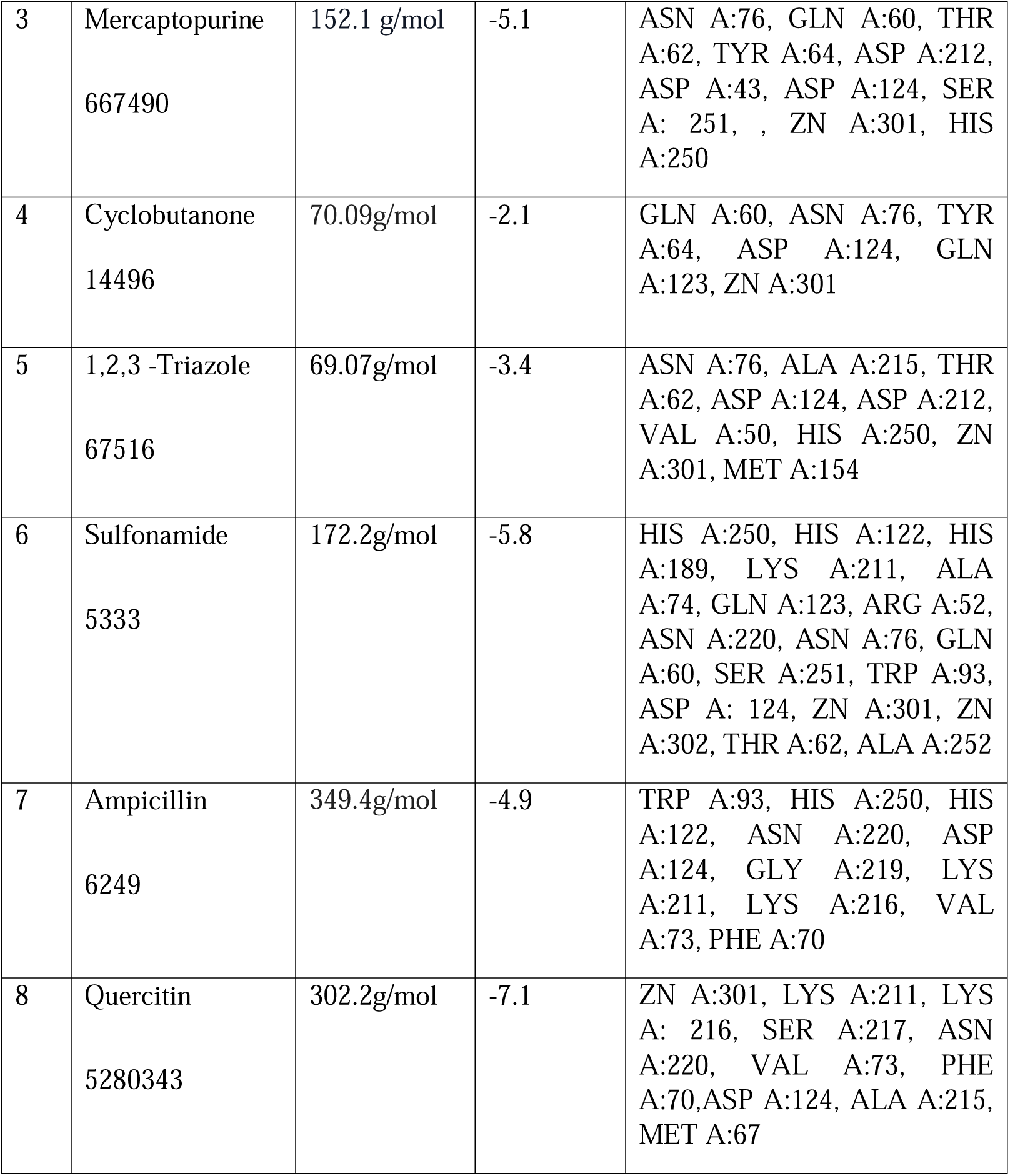
Docking results of standard ligands.

| Sl. No. | Name (pubchem CID) | Molecular weight | Binding Energy Kcal/mol | Residues interacted |
| --- | --- | --- | --- | --- |
| 1. | Penicillin<br>2349 | 334.4 g/mol | -7.1 | LYS A:211, ASN A: 220, ASN A:76, HIS A:250, HIS A:122, HIS A:189, SER A:217, SER A:251, ZN A:301, ZN A:301, ZN A:302, VAL A:73 |
| 2 | Benzylpenicillin<br>5904 | 334.4 g/mol | -8.0 | GLY A:219, ALA A:215, LYS A:211, HIS A:250, HIS A:122, HIS A:189, ZN A:301, ZN A:302, VAL A:73 |
| 3 | Mercaptopurine<br>667490 | 152.1 g/mol | -5.1 | ASN A:76, GLN A:60, THR A:62, TYR A:64, ASP A:212, ASP A:43, ASP A:124, SER A: 251, , ZN A:301, HIS A:250 |
| 4 | Cyclobutanone<br>14496 | 70.09g/mol | -2.1 | GLN A:60, ASN A:76, TYR A:64, ASP A:124, GLN A:123, ZN A:301 |
| 5 | 1,2,3 -Triazole<br>67516 | 69.07g/mol | -3.4 | ASN A:76, ALA A:215, THR A:62, ASP A:124, ASP A:212, VAL A:50, HIS A:250, ZN A:301, MET A:154 |
| 6 | Sulfonamide<br>5333 | 172.2g/mol | -5.8 | HIS A:250, HIS A:122, HIS A:189, LYS A:211, ALA A:74, GLN A:123, ARG A:52, ASN A:220, ASN A:76, GLN A:60, SER A:251, TRP A:93, ASP A: 124, ZN A:301, ZN A:302, THR A:62, ALA A:252 |
| 7 | Ampicillin<br>6249 | 349.4g/mol | -4.9 | TRP A:93, HIS A:250, HIS A:122, ASN A:220, ASP A:124, GLY A:219, LYS A:211, LYS A:216, VAL A:73, PHE A:70 |
| 8 | Quercitin<br>5280343 | 302.2g/mol | -7.1 | ZN A:301, LYS A:211, LYS A: 216, SER A:217, ASN A:220, VAL A:73, PHE A:70,ASP A:124, ALA A:215, MET A:67 |

The results obtained confirmed our suspision that sulfonamide could inhibit NDM-1 as it had a good binding affinity towards the protein. Thus, Sulfonamide derivatives were retrieved for further studies. The retrieved SDF file with ∼1400 sulfonamide compounds was screened against the identified active site of NDM-1. Top 5% compounds with respect to binding affinity were selected as hits from this for further studies.

### 2.3 ADMET STUDIES

ADMET (Adsorption, Distribution, Metabolism, Excretion and toxicity) properties of the top sulfonamide hits were to be studied for selecting druggable compounds. ADMETLab2.0 was used to carry out ADMET studies and Protox – II server was also used to extensively study toxicity of these compounds. Top 2 compounds (Pubchem CID: 118156306 and 46175386) among the hits found to be satisfying all the required properties were taken for simulation studies^14–15^.

### 2.4 MOLECULAR DYNAMICS SIMULATIONS

MD simulations were performed for the complexes the 2 hits formed with the target site in the NDM-1 protein to evaluate the stability of the complexes. GROMACSv5.0.4 was used for this study which operates based on linux commands. The complexes formed by 2 hit compounds with NDM-1 protein was retrieved in pdb format. Topology of these complexes were obtained using commands on gromacs followed by neutralizing and solvating the complexes by addition of a waterbox around the complexes. Equilibriation of the complexes was carried out to optimize the solute with solvent, first with constant number of atoms, volume and temperature (NVT) simulation followed by simulation where pressure and temperature were constant while volume was allowed to fluctuate (NPT). The systems were then subjected to 100 ns production MD run and the trajectories were plotted using ms excel to analyse the results^16–18^.

## 3. RESULTS AND DISCUSSION

### 3.1 PROTEIN PROFILE

The protein NDM-1 (PDB id: 6ny7) has [(5,7-dibromo-2-oxo-1,2-dihydroquinolin-4-yl methyl] phosphonic acid (L8J) as native ligand in it.

**Table 2:**
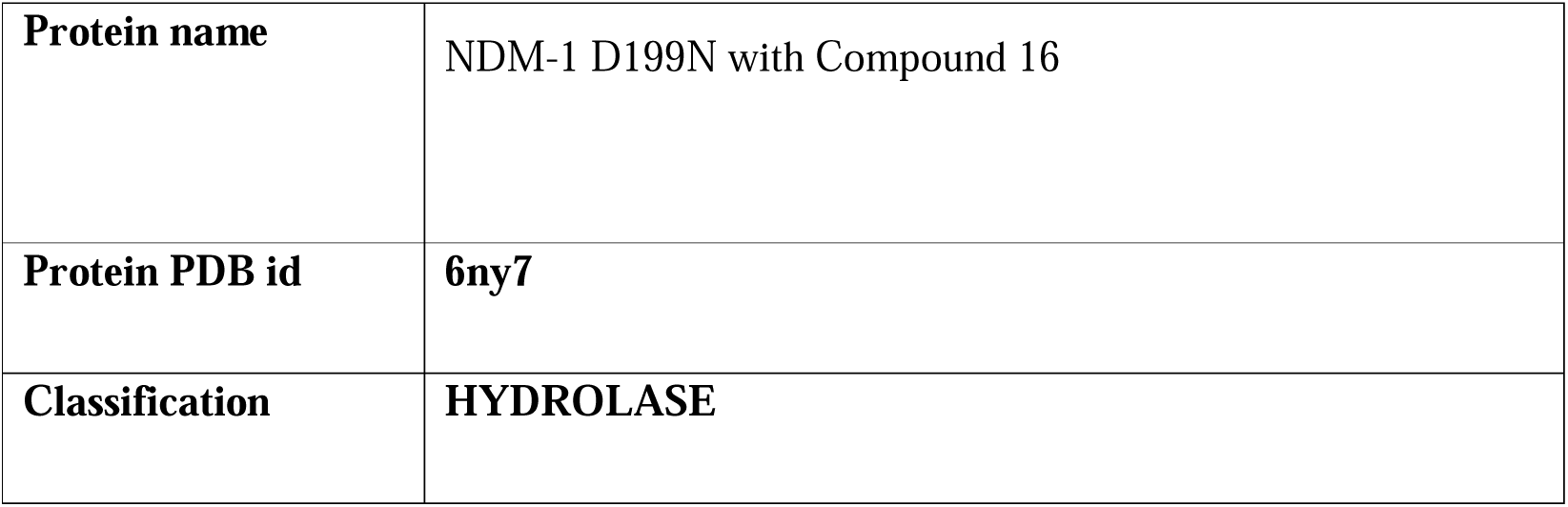
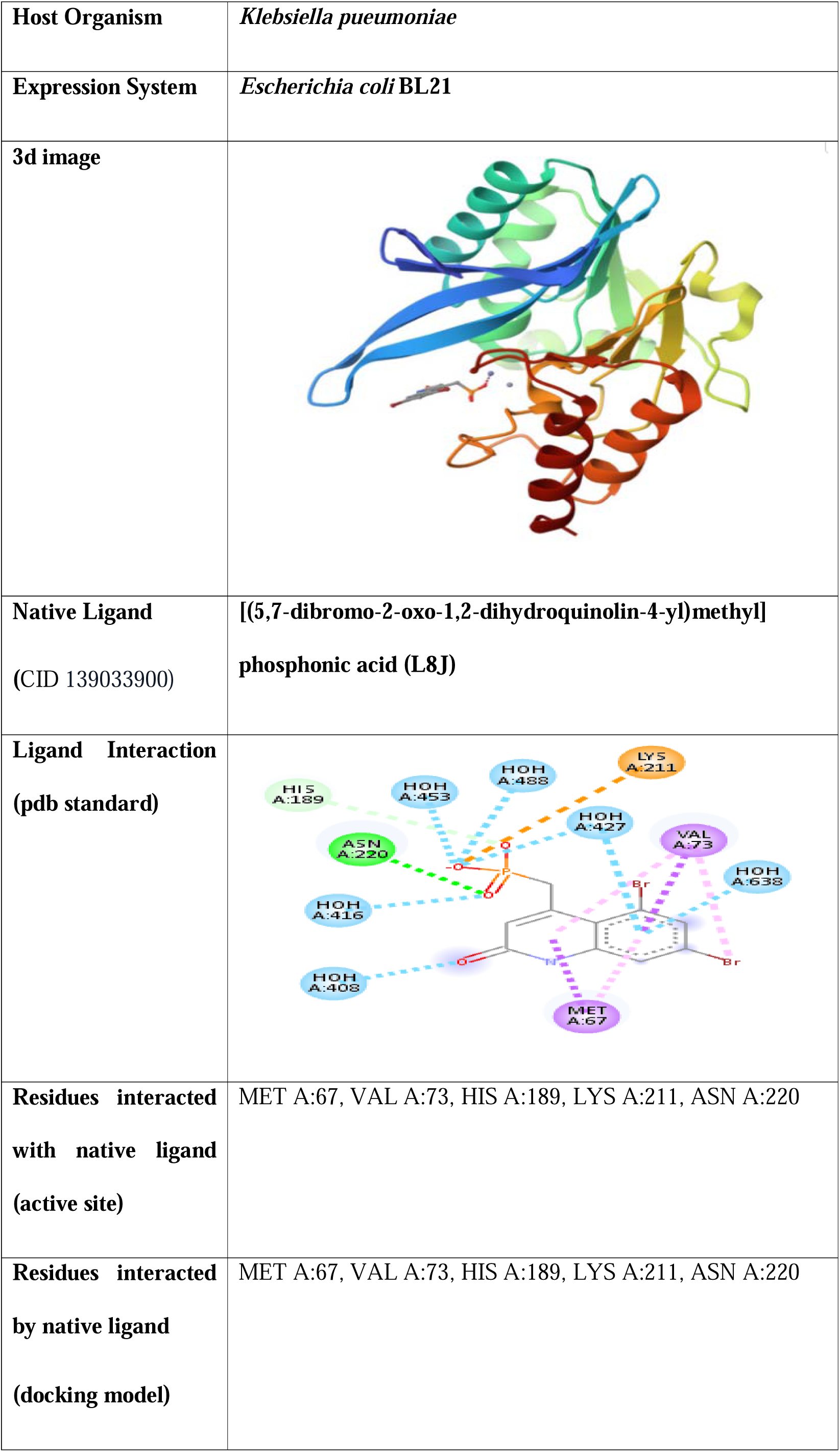
Protein Profile.

|  |  |
| --- | --- |
| **Protein name** | NDM-1 D199N with Compound 16 |
| **Protein PDB id** | **6ny7** |
| **Classification** | **HYDROLASE** |
| **Host Organism** | *Klebsiella pueumoniae* |
| **Expression System** | *Escherichia coli* BL21 |
| **3d image** |  |
| **Native Ligand**<br>(CID 139033900) | [(5,7-dibromo-2-oxo-1,2-dihydroquinolin-4-yl)methyl]<br>phosphonic acid (L8J) |
| **Ligand Interaction**<br>(pdb standard) |  |
| **Residues interacted with native ligand**<br>(active site) | MET A:67, VAL A:73, HIS A:189, LYS A:211, ASN A:220 |
| **Residues interacted by native ligand**<br>(docking model) | MET A:67, VAL A:73, HIS A:189, LYS A:211, ASN A:220 |

Docking of the native ligand **[(5,7-dibromo-2-oxo-1,2-dihydroquinolin-4-yl) methyl] phosphonic acid** with NDM-1 active site was performed. The interactions of native ligand with NDM-1 protein (PDB id: 6ny7) obtained from PDB database were compared with docking results during validation of docking model. Both showed the same interactions which validated our docking model.

### 3.2 DOCKING STUDIES

About 1lakh sulfonamide derivative compounds were found in pubchem database. By applying Verber’s and Lipinski’s rules of drug likeliness, the number of compounds were narrowed down to ∼1400 which were downloaded and screened along with the native ligand, positive control and negative control against the active site of NDM-1 protein (6ny7) using Autodock Vina in PyRx. Top 60 (5% of the 1400 compounds screened) compounds with respect to binding energy were selected as best hits and their details are given in the following tables.

**Table 3:** Virtual Screening results of best hits.

| Compound (Pubchem CID) | IUPAC Name | Binding Energy<br>(Kcal/mol) |
| --- | --- | --- |
| 103646811 | (3S)-3-hydroxypyrrolidine-1-sulfonamide | -5.1 |
| **118156306** | (3Z)-N-hydroxypenta-1,3-diene-2-sulfonamide | -5.4 |
| 156787384 | 4-fluoro-1H-pyrazole-5-sulfonamide | -5.1 |
| 136574701 | 5-methyl-1H-imidazole-4-sulfonamide | -5.1 |
| **139364956** | (1R,2R)-2-hydroxycyclopentane-1-sulfonamide | -5.4 |
| 139388953 | 4-methyl-1H-pyrazole-5-sulfonamide | -5.2 |
| 64514083 | 3-methylcyclopentane-1-sulfonamide | -5.2 |
| 139389384 | 5-methyl-1H-pyrazole-3-sulfonamide | -5.1 |
| 141764153 | 2-hydroxypyrrolidine-1-sulfonamide | -5.4 |
| 156030339 | (1R,2R)-2-methylcyclopentane-1-sulfonamide | -5.3 |
| 156530860 | 2-fluorothiophene-3-sulfonamide | -5.1 |
| 45080568 | 2-oxo-1,3-oxazolidine-3-sulfonamide | -5.3 |
| 46175386 | N-hydroxyfuran-2-sulfonamide | -5.1 |
| 65050329 | 2-methylcyclopentane-1-sulfonamide | -5.2 |
| 67028586 | 2-oxo-1,3-oxazole-3-sulfonamide | -5.1 |
| 103646815 | (3R)-3-hydroxypyrrolidine-1-sulfonamide | -5.2 |
| 17976419 | Cyclohexene sulfonamide | -5.3 |
| 7370 | Benzenesulfonamide | -5.2 |
| 75536 | Cyclohexanesulfonamide | -5.2 |
| 90142128 | 3-methylfuran-2-sulfonamide | -5.2 |

### 3.3 ADMET STUDIES

After studying the results of virtual screening, Top 60 (5% of the 1400 compounds screened) compounds with respect to binding energy were selected as best hits for conducting ADMET studies which was carried out using online servers, ADMETLab2.0 and Pro Tox-II. The results obtained from ADMET studies are shown below. Among these compounds, 26 were selected for further studies as they had better ADME profile and were non-toxic. The results obtained from ADMET studies are shown below. Among these compounds, 26 were selected for further studies as they had better ADME profile and were non-toxic.

**Table 4:** Toxicity prediction results of best hits.

| **Compound**<br>**(Pubchem CID)** | **Hepatotoxicity** | **Carcinogenicity** | **Immunotoxicity** | **Mutagenicity** | **Cytotoxicity** |
| --- | --- | --- | --- | --- | --- |
| **103646811** | **Inactive** | **Inactive** | **Inactive** | **Inactive** | **Inactive** |
|  | **0.22** | **0.48** | **0.01** | **0.34** | **0.36** |
| **75536** | **Inactive** | **Inactive** | **Inactive** | **Inactive** | **Inactive** |
|  | **0.16** | **0.44** | **0.01** | **0.17** | **0.23** |
| **7370** | **Inactive** | **Inactive** | **Inactive** | **Inactive** | **Inactive** |
|  | **0.16** | **0.46** | **0.01** | **0.17** | **0.23** |
| **139364956** | **Inactive** | **Inactive** | **Inactive** | **Inactive** | **Inactive** |
|  | **0.26** | **0.39** | **0.01** | **0.30** | **0.27** |
| **141764153** | **Inactive** | **Inactive** | **Inactive** | **Inactive** | **Inactive** |
|  | **0.27** | **0.41** | **0.01** | **0.33** | **0.34** |
| **152760603** | **Inactive** | **Inactive** | **Inactive** | **Inactive** | **Inactive** |
|  | **0.27** | **0.47** | **0.01** | **0.31** | **0.44** |
| **156030339** | **Inactive** | **Inactive** | **Inactive** | **Inactive** | **Inactive** |
|  | **0.26** | **0.47** | **0.01** | **0.29** | **0.25** |
| **156530860** | **Inactive** | **Inactive** | **Inactive** | **Inactive** | **Inactive** |
|  | **0.25** | **0.37** | **0.02** | **0.27** | **0.22** |
| **17976419** | **Inactive** | **Inactive** | **Inactive** | **Inactive** | **Inactive** |
|  | **0.25** | **0.48** | **0.01** | **0.30** | **0.29** |
| **45080568** | **Inactive** | **Inactive** | **Inactive** | **Inactive** | **Inactive** |
|  | **0.48** | **0.44** | **0.05** | **0.32** | **0.39** |
| **64514083** | **Inactive** | **Inactive** | **Inactive** | **Inactive** | **Inactive** |
|  | **0.26** | **0.48** | **0.01** | **0.29** | **0.25** |
| **65050329** | **Inactive** | **Inactive** | **Inactive** | **Inactive** | **Inactive** |
| **118910321** | **Inactive** | **Inactive** | **Inactive** | **Inactive** | **Inactive** |
|  | **0.26** | **0.47** | **0.01** | **0.29** | **0.25** |
|  | **0.22** | **0.48** | **0.01** | **0.35** | **0.25** |
| **67028586** | **Inactive** | **Inactive** | **Inactive** | **Inactive** | **Inactive** |
| **46175386** | **Inactive** | **Inactive** | **Inactive** | **Inactive** | **Inactive** |
|  | **0.48** | **0.44** | **0.05** | **0.32** | **0.39** |
| **103646815** | **Inactive** | **Inactive** | **Inactive** | **Inactive** | **Inactive** |
|  | **0.22** | **0.48** | **0.01** | **0.34** | **0.36** |
| **90142128** | **Inactive** | **Inactive** | **Inactive** | **Inactive** | **Inactive** |
|  | **0.24** | **0.45** | **0.001** | **0.28** | **0.23** |
| **118156306** | **Inactive** | **Inactive** | **Inactive** | **Inactive** | **Inactive** |
|  | **0.36** | **0.48** | **0.01** | **0.50** | **0.29** |
|  | **0.42** | **0.48** | **0.01** | **0.43** | **0.34** |

**(The values in table denote the probability of the specific toxicity occurring on administration of the compound as a drug. Compounds getting the result inactive with a probability of less than 0.49 as considered as hits)**

**Table 5:**
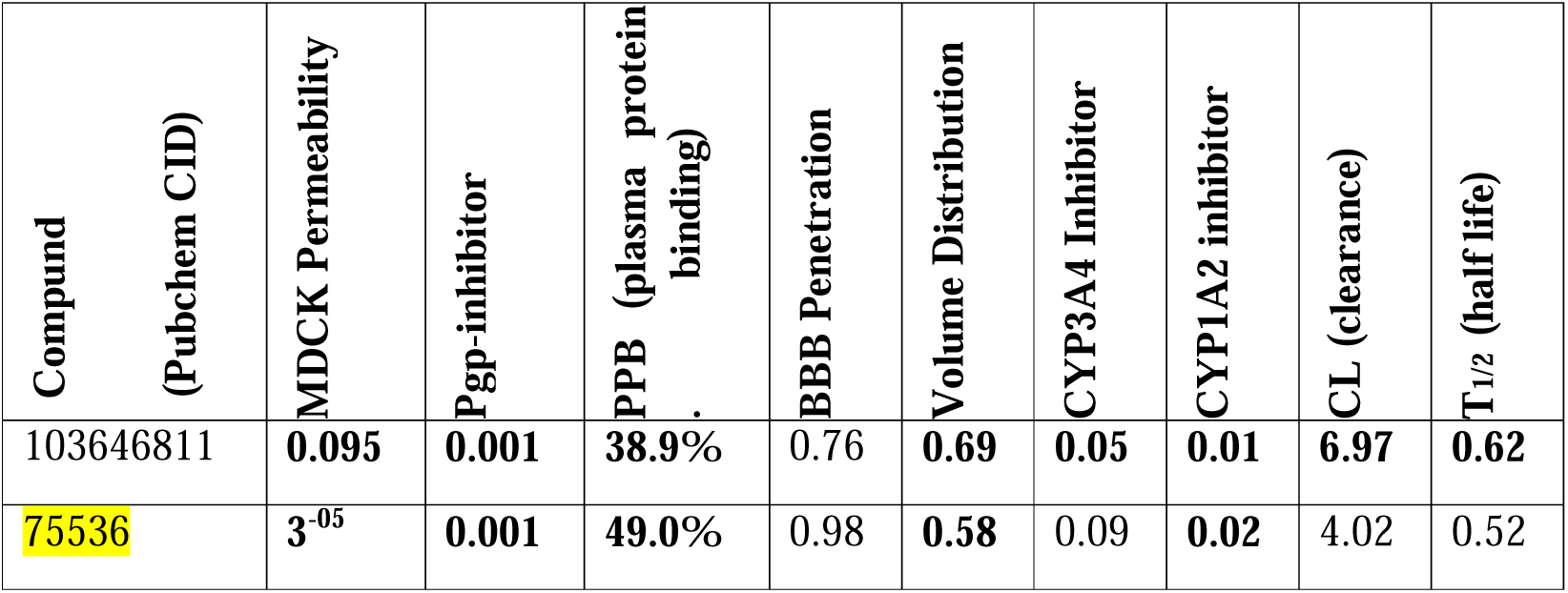
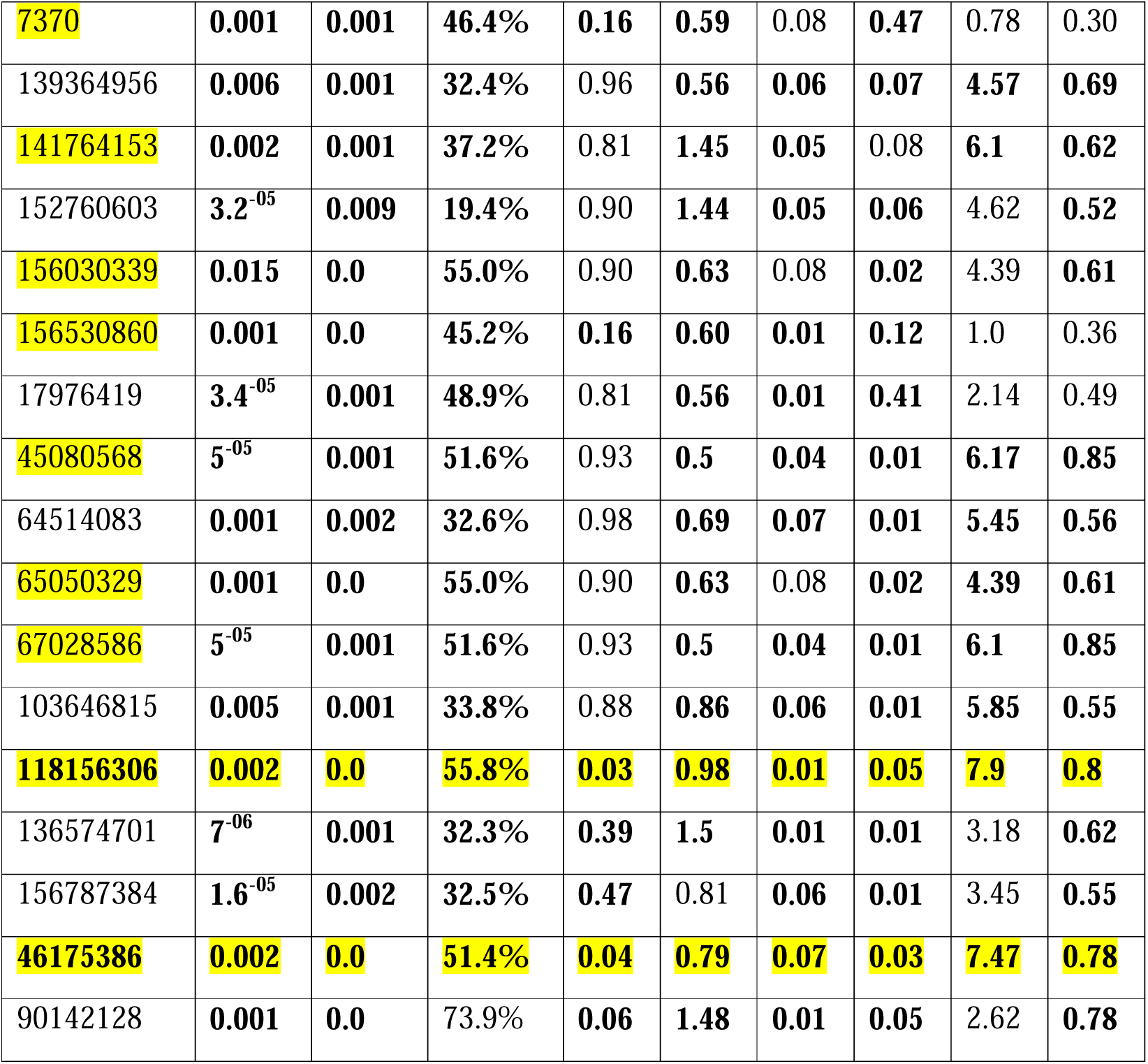
ADME Prediction results of best hits.

| **Compound**<br>**(Pubchem CID)** | **MDCK Permeability** | **Pgp-inhibitor** | **PPB (plasma protein binding)** | **BBB Penetration** | **Volume Distribution** | **CYP3A4 Inhibitor** | **CYP1A2 inhibitor** | **CL (clearance)** | **T<sub>1/2</sub> (half life)** |
| --- | --- | --- | --- | --- | --- | --- | --- | --- | --- |
| 103646811 | **0.095** | **0.001** | **38.9%** | 0.76 | **0.69** | **0.05** | **0.01** | **6.97** | **0.62** |
| 75536 | **3<sup>-05</sup>** | **0.001** | **49.0%** | 0.98 | **0.58** | 0.09 | **0.02** | 4.02 | 0.52 |
| 7370 | 0.001 | 0.001 | 46.4% | 0.16 | 0.59 | 0.08 | 0.47 | 0.78 | 0.30 |
| 139364956 | 0.006 | 0.001 | 32.4% | 0.96 | 0.56 | 0.06 | 0.07 | 4.57 | 0.69 |
| 141764153 | 0.002 | 0.001 | 37.2% | 0.81 | 1.45 | 0.05 | 0.08 | 6.1 | 0.62 |
| 152760603 | 3.2<sup>-05</sup> | 0.009 | 19.4% | 0.90 | 1.44 | 0.05 | 0.06 | 4.62 | 0.52 |
| 156030339 | 0.015 | 0.0 | 55.0% | 0.90 | 0.63 | 0.08 | 0.02 | 4.39 | 0.61 |
| 156530860 | 0.001 | 0.0 | 45.2% | 0.16 | 0.60 | 0.01 | 0.12 | 1.0 | 0.36 |
| 17976419 | 3.4<sup>-05</sup> | 0.001 | 48.9% | 0.81 | 0.56 | 0.01 | 0.41 | 2.14 | 0.49 |
| 45080568 | 5<sup>-05</sup> | 0.001 | 51.6% | 0.93 | 0.5 | 0.04 | 0.01 | 6.17 | 0.85 |
| 64514083 | 0.001 | 0.002 | 32.6% | 0.98 | 0.69 | 0.07 | 0.01 | 5.45 | 0.56 |
| 65050329 | 0.001 | 0.0 | 55.0% | 0.90 | 0.63 | 0.08 | 0.02 | 4.39 | 0.61 |
| 67028586 | 5<sup>-05</sup> | 0.001 | 51.6% | 0.93 | 0.5 | 0.04 | 0.01 | 6.1 | 0.85 |
| 103646815 | 0.005 | 0.001 | 33.8% | 0.88 | 0.86 | 0.06 | 0.01 | 5.85 | 0.55 |
| 118156306 | 0.002 | 0.0 | 55.8% | 0.03 | 0.98 | 0.01 | 0.05 | 7.9 | 0.8 |
| 136574701 | 7<sup>-06</sup> | 0.001 | 32.3% | 0.39 | 1.5 | 0.01 | 0.01 | 3.18 | 0.62 |
| 156787384 | 1.6<sup>-05</sup> | 0.002 | 32.5% | 0.47 | 0.81 | 0.06 | 0.01 | 3.45 | 0.55 |
| 46175386 | 0.002 | 0.0 | 51.4% | 0.04 | 0.79 | 0.07 | 0.03 | 7.47 | 0.78 |
| 90142128 | 0.001 | 0.0 | 73.9% | 0.06 | 1.48 | 0.01 | 0.05 | 2.62 | 0.78 |

### Ranges Desciption

( MDCK Permeability: < 2 * 10^-6^ cm/s; Pgp inhibitor:0 = non inhibitor and 1 = inhibitor; PPB: < 90%; BBB Penetration: 0 = BBB-and 1 = BBB+; Volume Distribution: 0.04-20L/kg; CYP3A4 inhibitor: 0 = non inhibitor and 1 = inhibitor; CYP1A2 inhibitor: 0 = non substrate and 1 = substrate; Clearance: Low = < 5 ml/min/kg, Moderate = 5-15 ml/min/kg, High = > 20 ml/min/kg; t _½_: 0 = short half life and 1 = long half life )

**Table 6:**
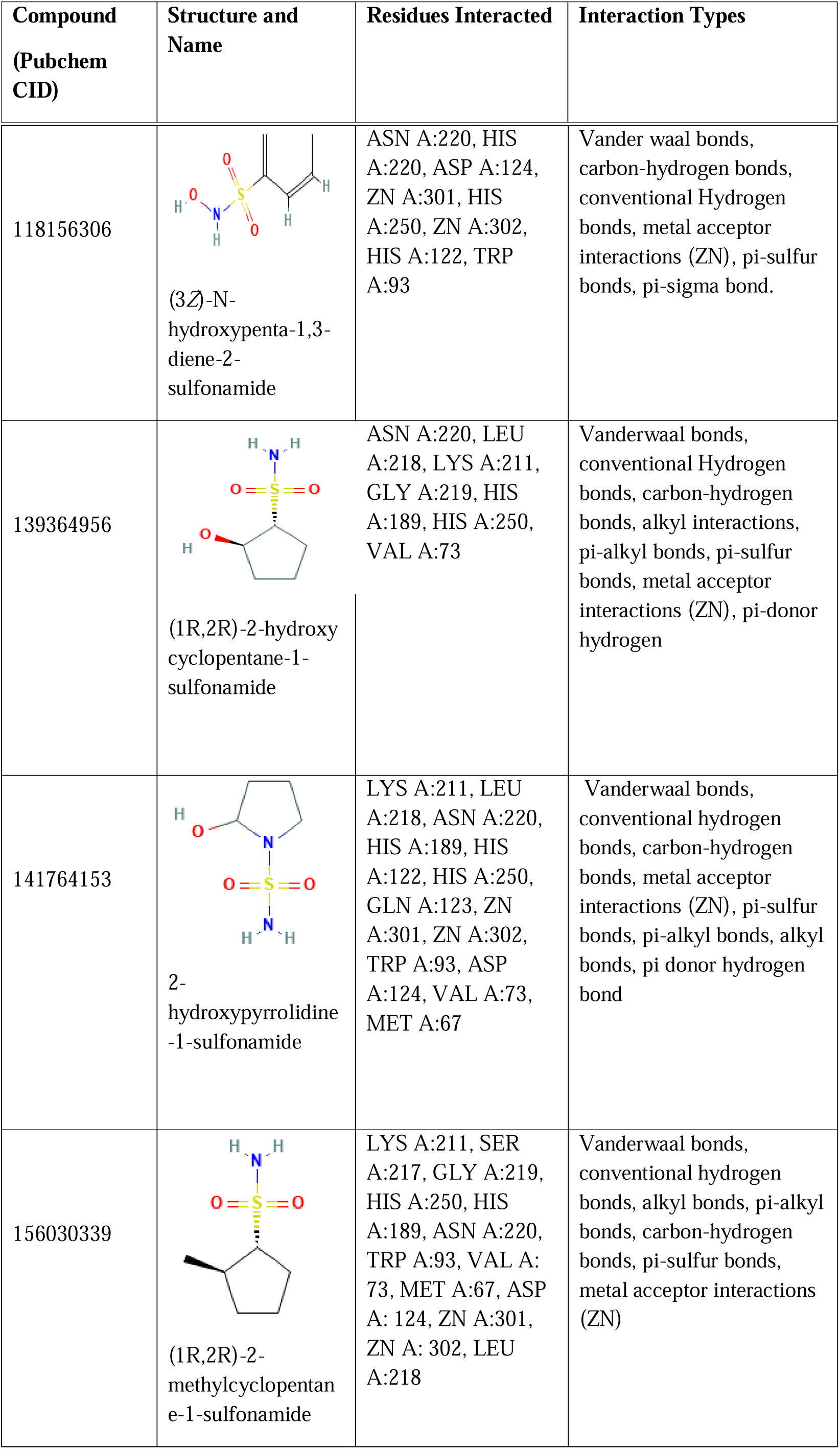
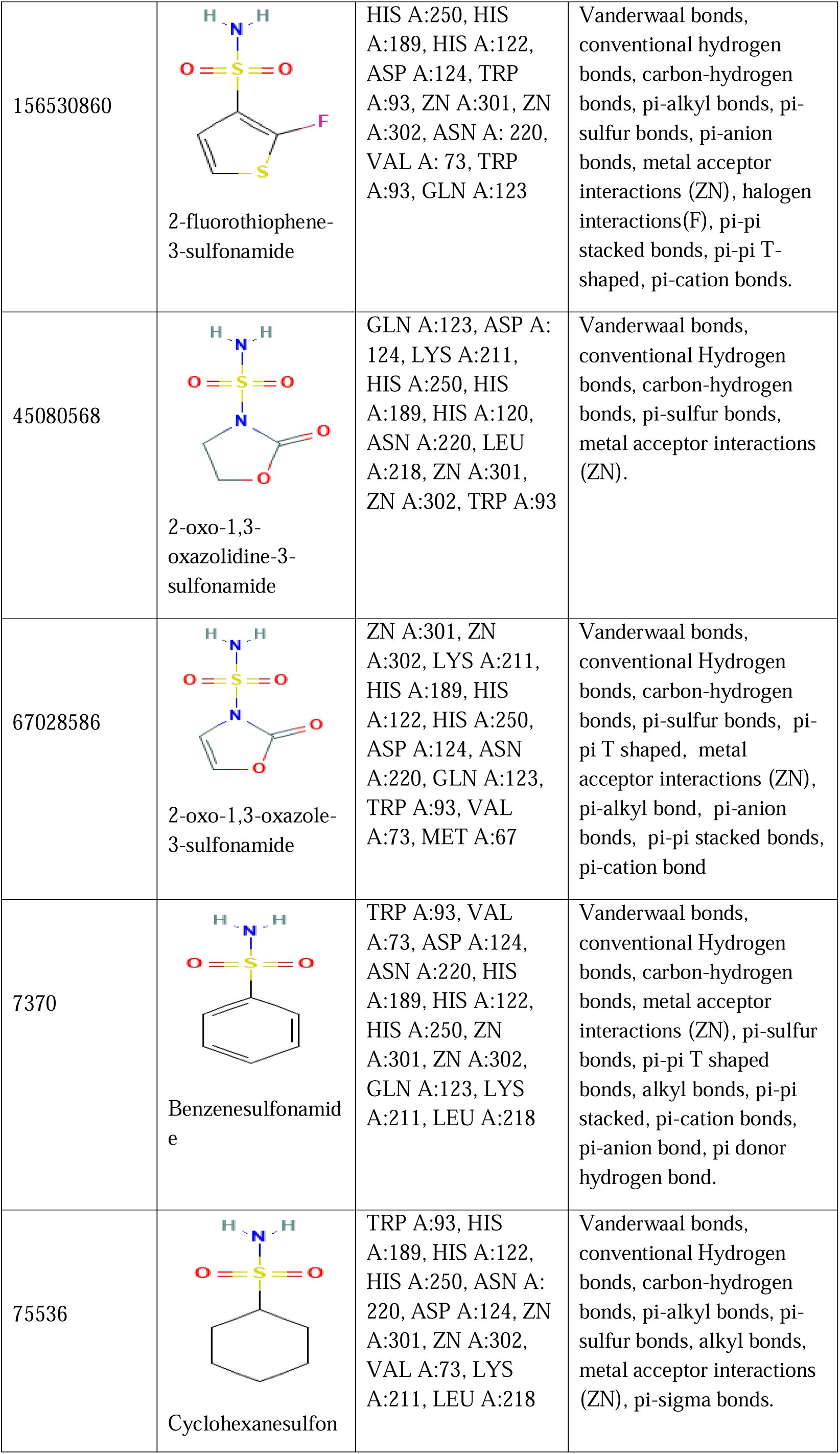
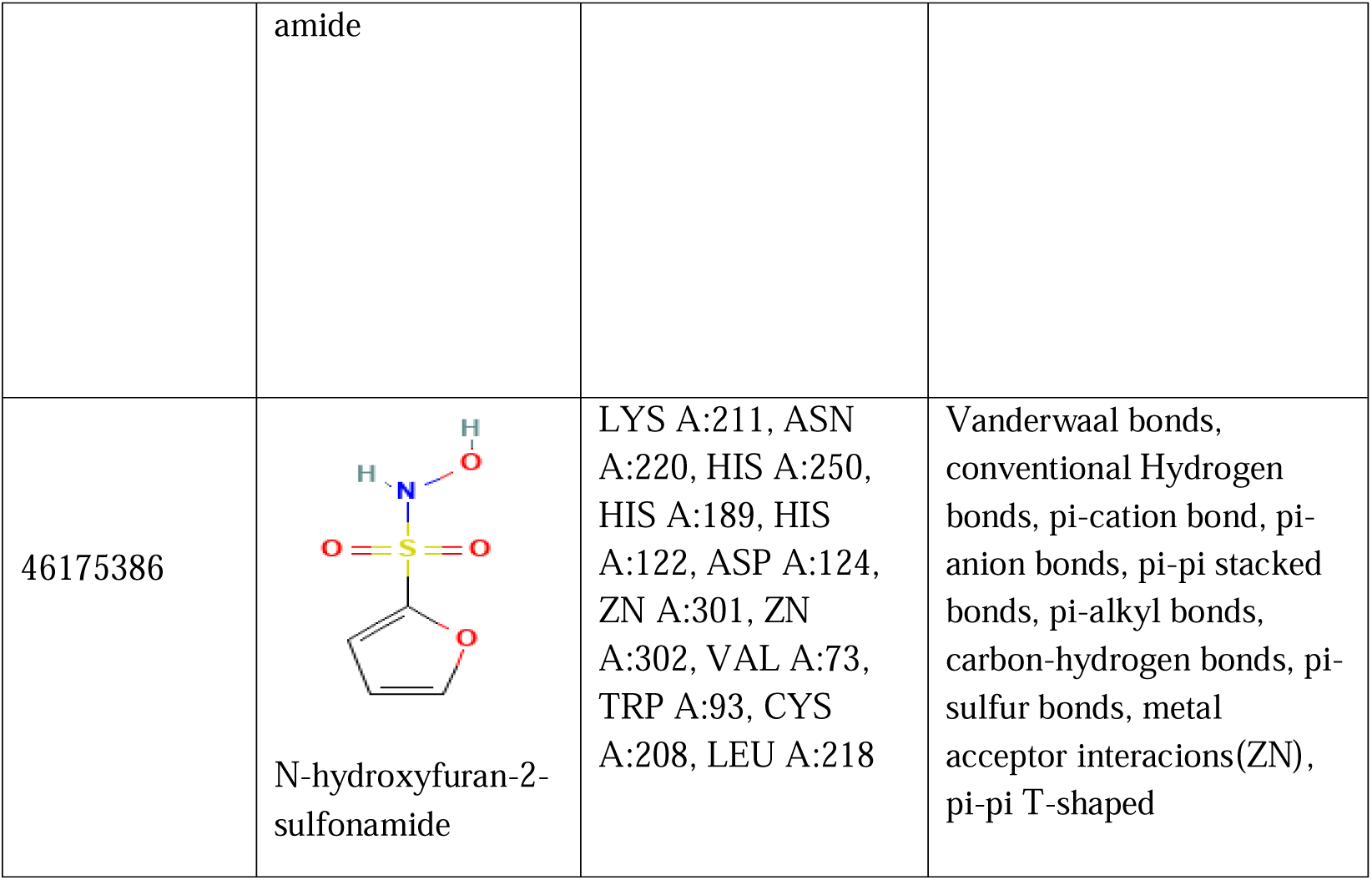
Molecular Profile and Interactions of Best hits.

2 Compounds with CID 46175386 and 118156306 were prioritized for molecular dynamics simulations owing to their favourable binding affinities and potential to inhibit NDM-1 via a metal chel tion mechanism, supported by the presence of Zn² interactions in their binding profiles.

**Fig 2:**
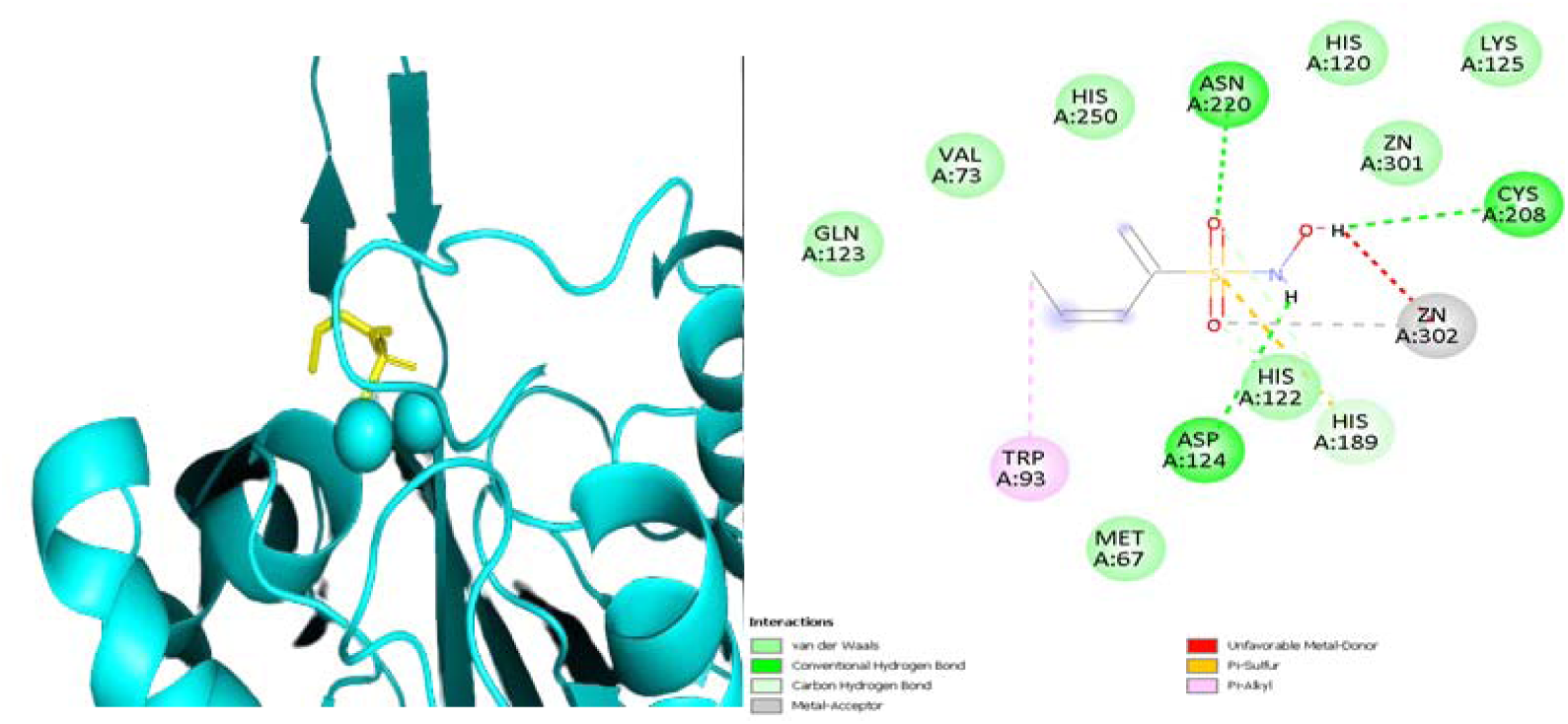
**Visual representation of Interactions of of N-hydroxyfuran-2-sulfonamide with NDM-1**

**Fig 3:**
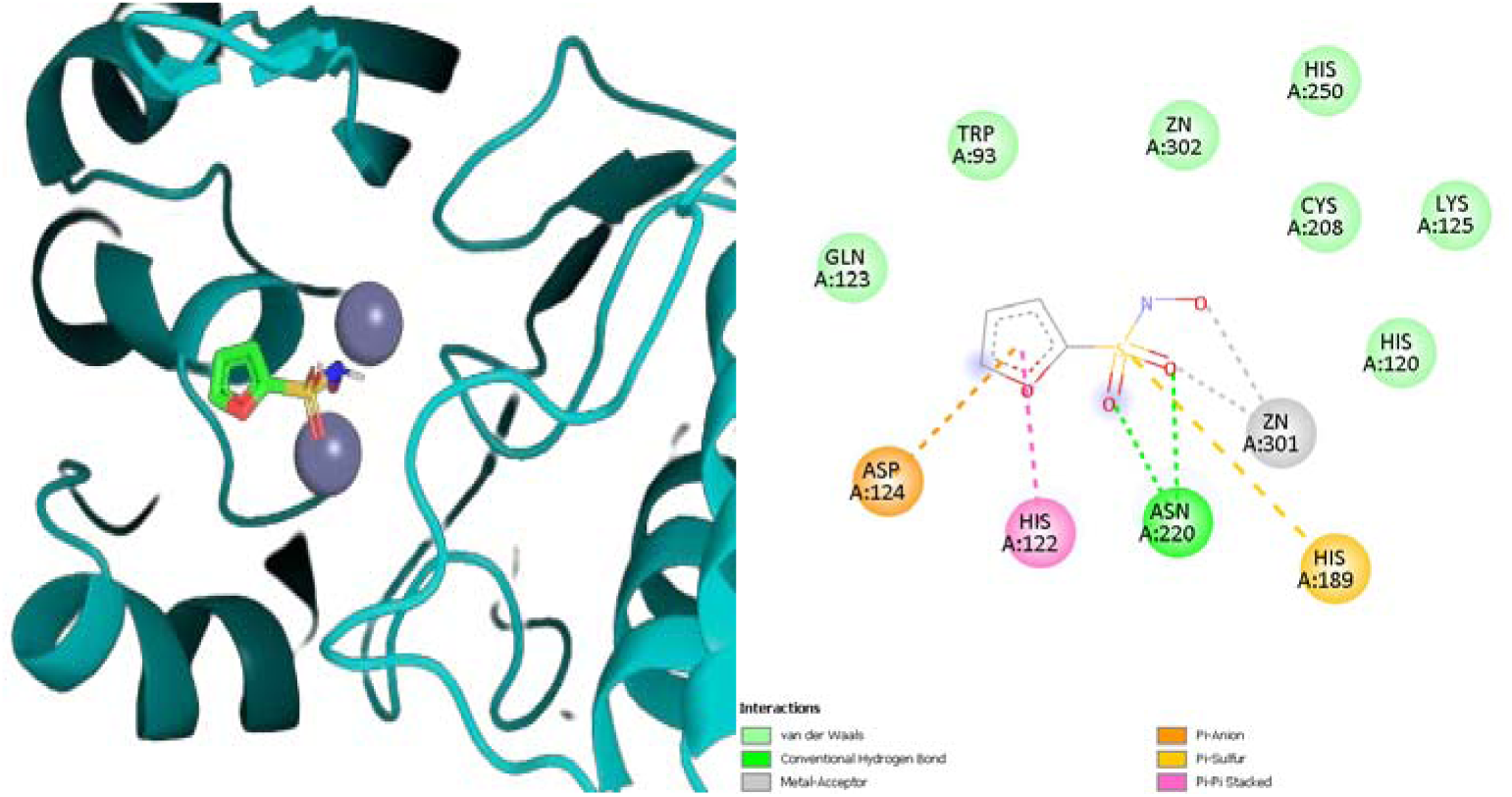
**Visual representation of interactions of (3Z)-N-hydroxypenta-1,3-diene-2-sulfonamide NDM-1**

### 3.4 MD SIMULATIONS

The chain to which ligand was bound contained 228 residues numbered from 42 to 270 and the other residues were deleted before simulation to reduce computational workload. The complexes formed by 2 hit compounds N-hydroxyfuran-2-sulfonamide (CID:46175386) and (3Z)-N-hydroxypenta-1,3-diene-2-sulfonamide (CID:118156306) with NDM-1 protein were subjected to MD simulations and the results were compared to md simulation of protein system alone (NDM-1 - control) to evaluate their stability an results are discussed below:

#### 3.4.1 NDM-1(3Z)-N-hydroxypenta-1,3-diene-2-sulfonamide complex(118156306) COMPLEX

Root mean square deviation is a parameter used to measure the difference between the backbone of the protein from its initial structure to its final position which enables us to evaluate the stability of the protein - ligand complexes.

The RMSD plot is seen going upwards initially suggesting early equilibration. There is a slight increase in RMSD value upto 10ns which represents an equilibration phase where the complex transitions into a more stable state with some fluctuation. RMSD is in between 0.1 to 0.25 throughout the simulation after i phase which is not a significant fluctuation. One peak is observed in the complex at 35ns compared to the protein(control) possibly induced by ligand binding. This shows that the complex is stable with minimal fluctuation after equilibration is achieved which should provide sufficient duration for the ligand to act on the protein.

**Fig 4.**
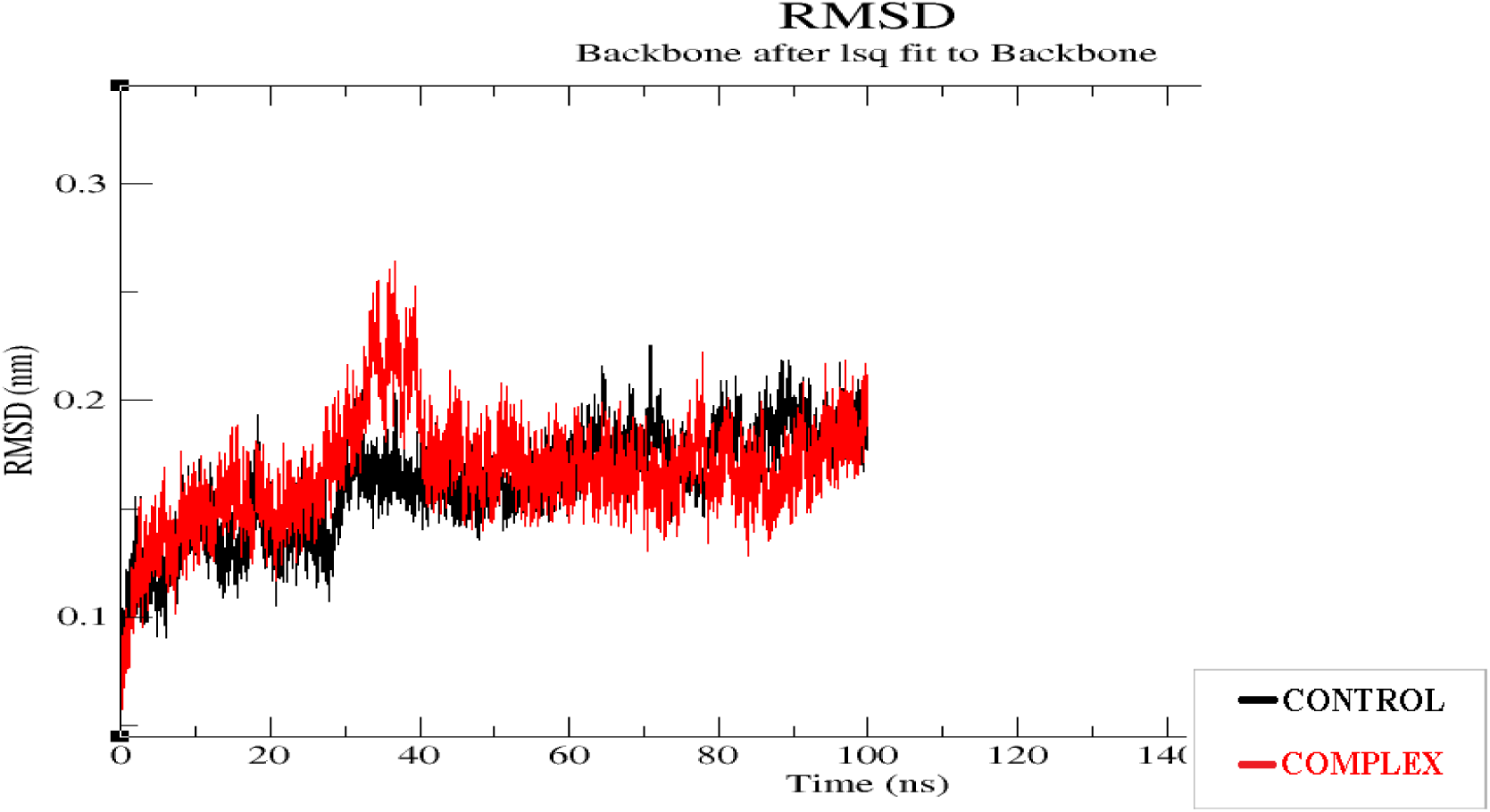
RMSD plot of NDM-1 and (3Z)-N-hydroxypenta-1,3-diene-2-sulfonamide complex.

**Fig 5.**
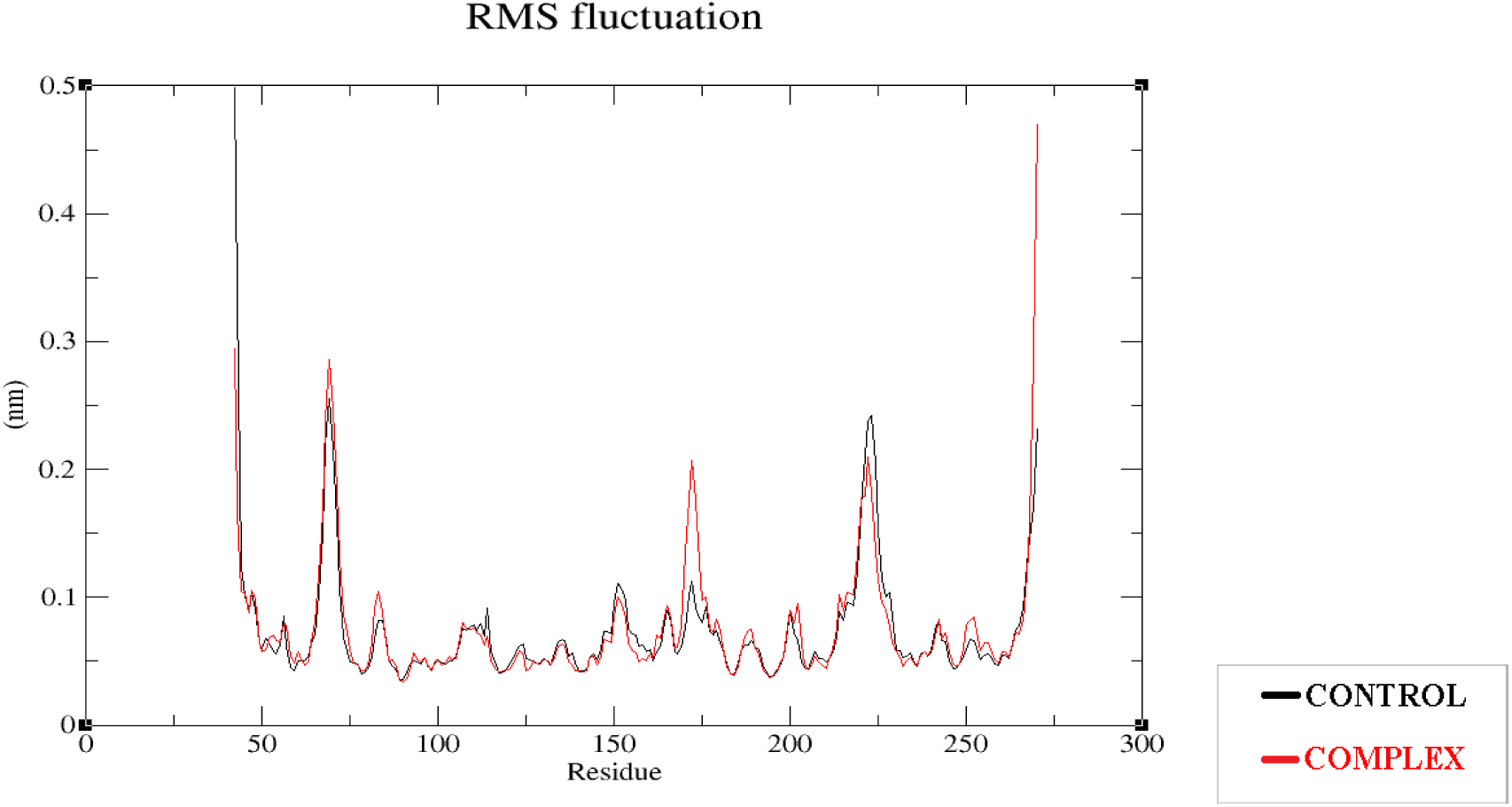
RMSF plot of NDM-1 and (3Z)-N-hydroxypenta-1,3-diene-2-sulfonamide complex. RMSF measures the average deviation of atomic positions from their mean position along the simulation. Some small fluctuations are seen at residues 67,73,189,220 probably as a dynamic response to the binding as all those residues are in the binding site. No significant fluctuation is exhibited in other regions indicative of structurally rigid areas. The data suggests that the protein retains its structural integrity with localised flexibility and no significant fluctuation has occurred due to the ligand binding to the protein throughout the 100ns of simulation.

**Fig 6.**
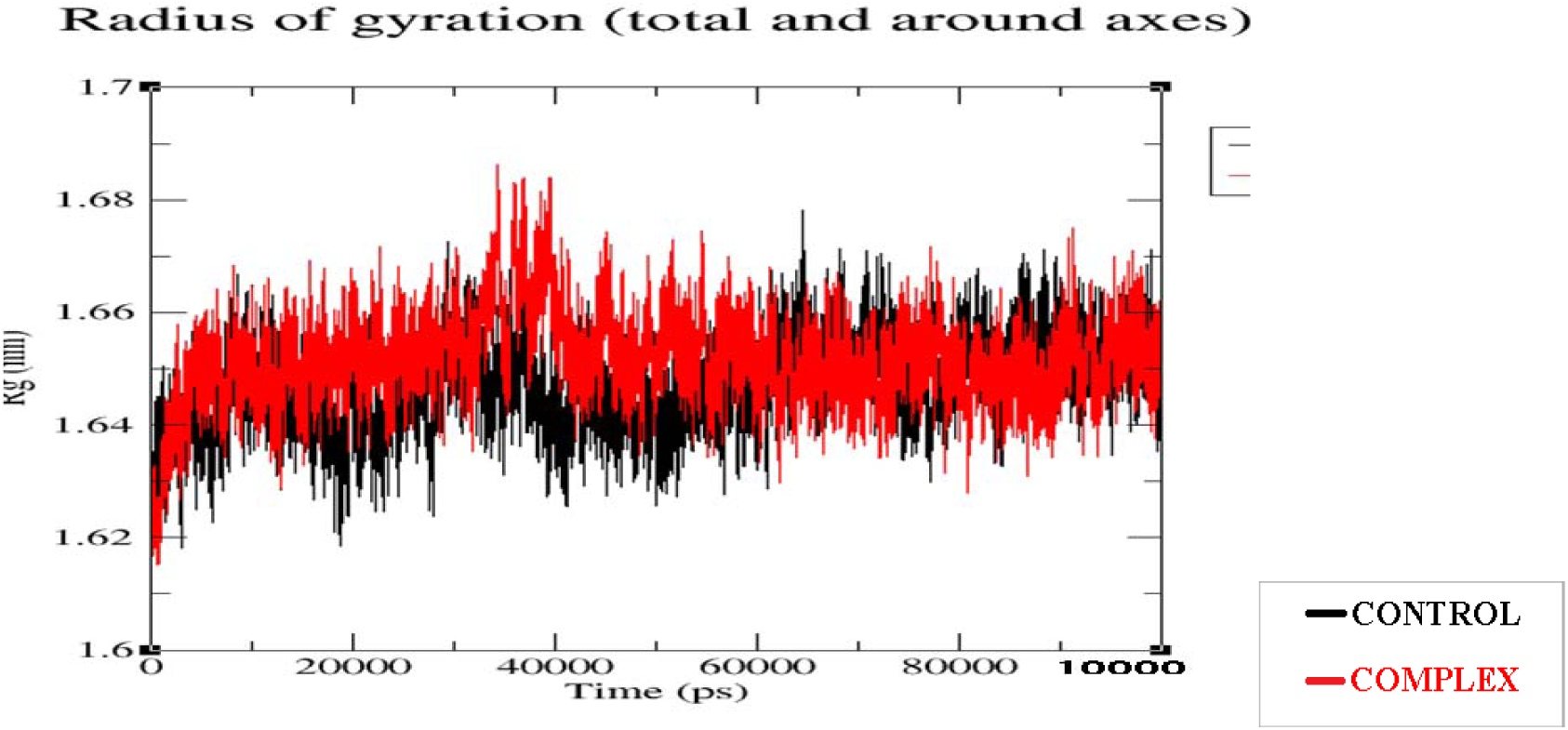
Rg plot of NDM-1 and (3Z)-N-hydroxypenta-1,3-diene-2-sulfonamide complex. Radius of gyration is an index to monitor the structural formation process used to assess compactne s of protein. There is a steady increase in Rg initially upto 1.64nm at 10000ps suggesting the adaptation of complex to a more compact confirmation from its initial confirmation. No significant fluctuation is seen afterwards suggesting that the complex retains its compactness throughout the simulation

**Fig 7.**
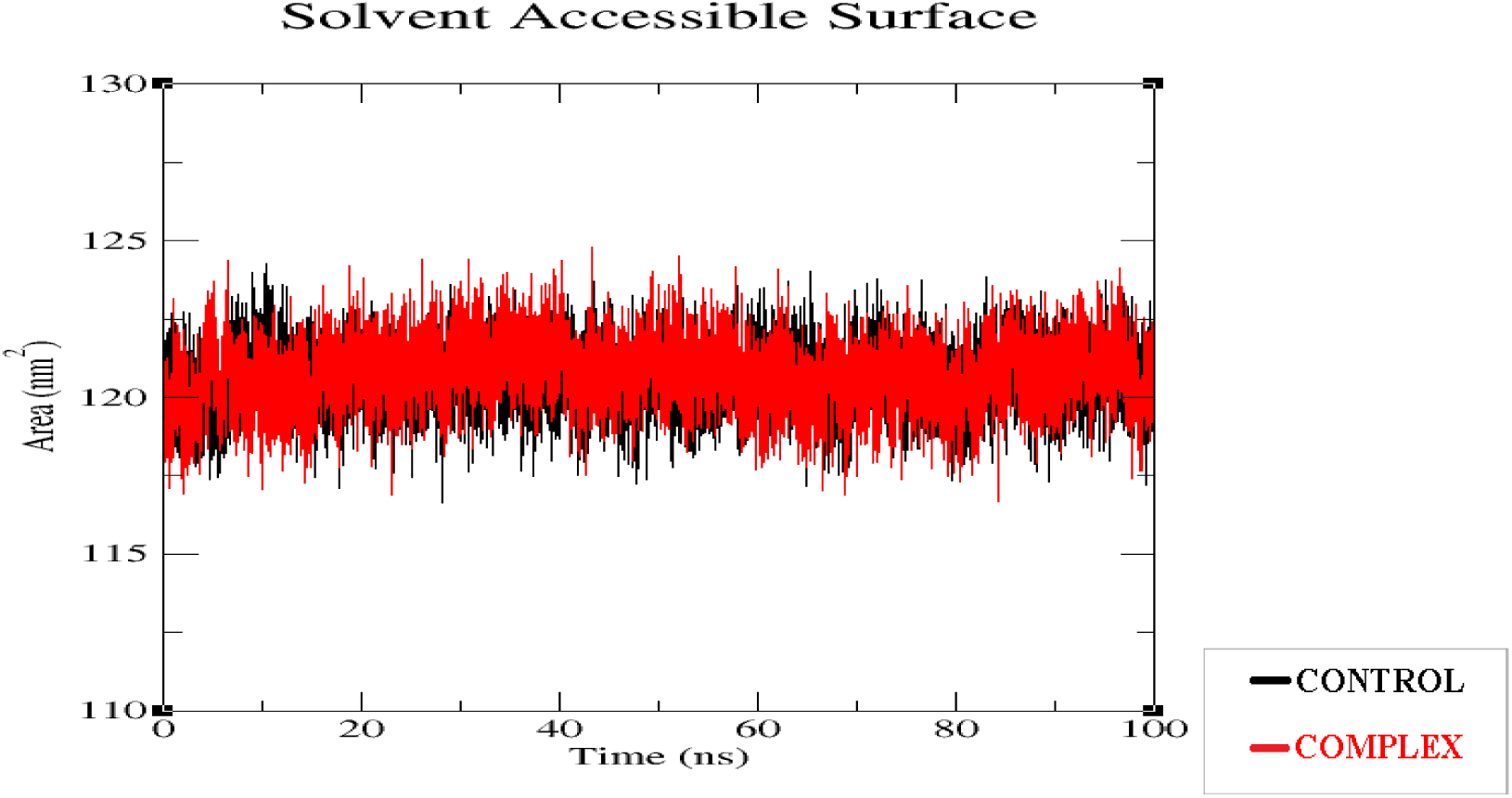
SASA plot of NDM-1 and (3Z)-N-hydroxypenta-1,3-diene-2-sulfonamide complex. Solvent Accessible Surface Area is a measure of the surface area of a molecule that is exposed to the solvent in a given system. SASA is often used as an energy term to describe the interactions between the molecule and the solvent. There is an initial fluctuation in SASA in both complex and the protein probably because of ligand binding and confirmational changes followed by a stable phase where SASA roughly remains around120nm^2^ in both control and complex throughout the simulation which indicates the complex doe not undergo major structural rearrangements for the remaining duration of simulation which means the complex is stable enough for the ligand to act on the protein.

#### 3.4.2 NDM1 – N-hydroxyfuran-2-sulfonamide (46175386) COMPLEX

**Fig 8.**
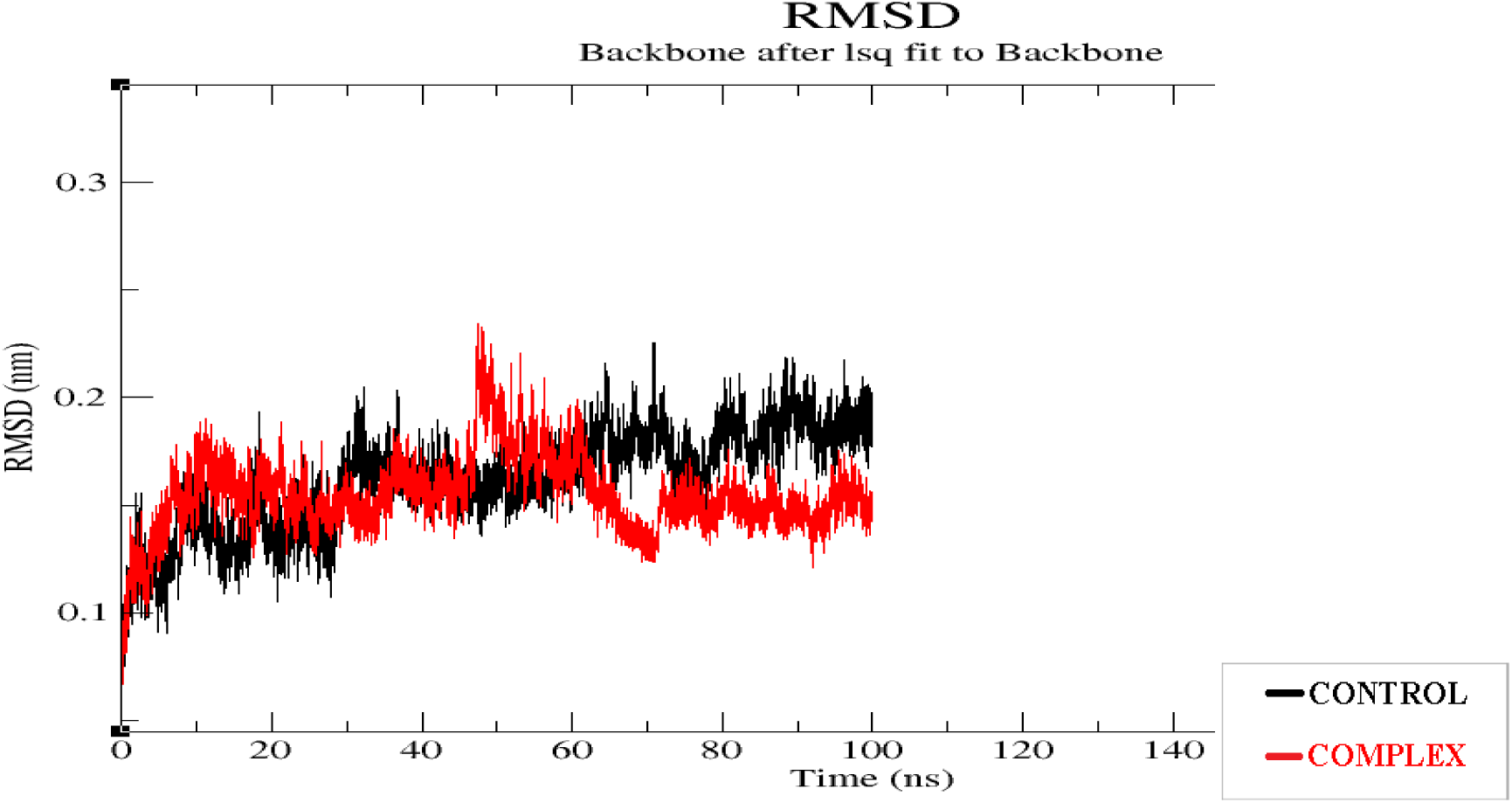
RMSD plot of NDM1 – N-hydroxyfuran-2-sulfonamide complex. The RMSD value is seen going upwards initially suggesting early equilibration followed by some small fluctuations upto 44ns where an upward fluctuatiom in RMSD is seen followed by a downward fluctuation due to the confirmational changes caused by ligand binding. The system appears to be stablized at 65ns where it has transitioned into a more stable state. RMSD ranges in between 0.1 to 0.22 throughout the simulation which is not a significant fluctuation. This indicates that the complex is stable with minimal fluctuation after equilibration is achieved which should provide sufficient duration for the ligand to act on the protein.

**Fig 9.**
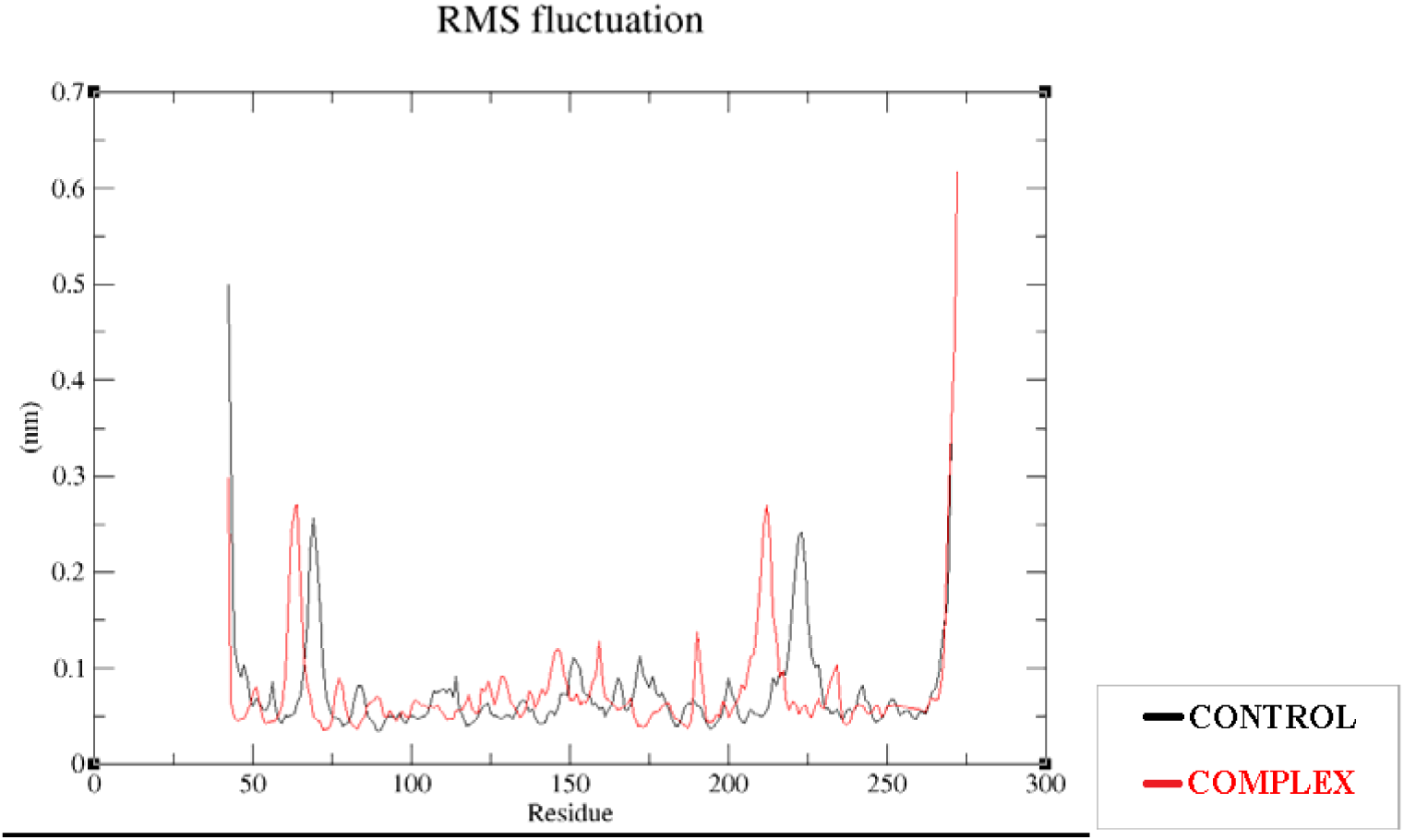
RMSF plot of NDM1 – N-hydroxyfuran-2-sulfonamide complex. There is some fluctuation in RMSF of the complex when compared to control at residues 67,73,189and 220 probably induced by ligand binding but none of the fluctuations are significantly high. This data suggests that the protein retains its structural integrity with localised flexibility and no significant change is caused due to ligand binding throughout the duration of simulation.

**Fig 10.**
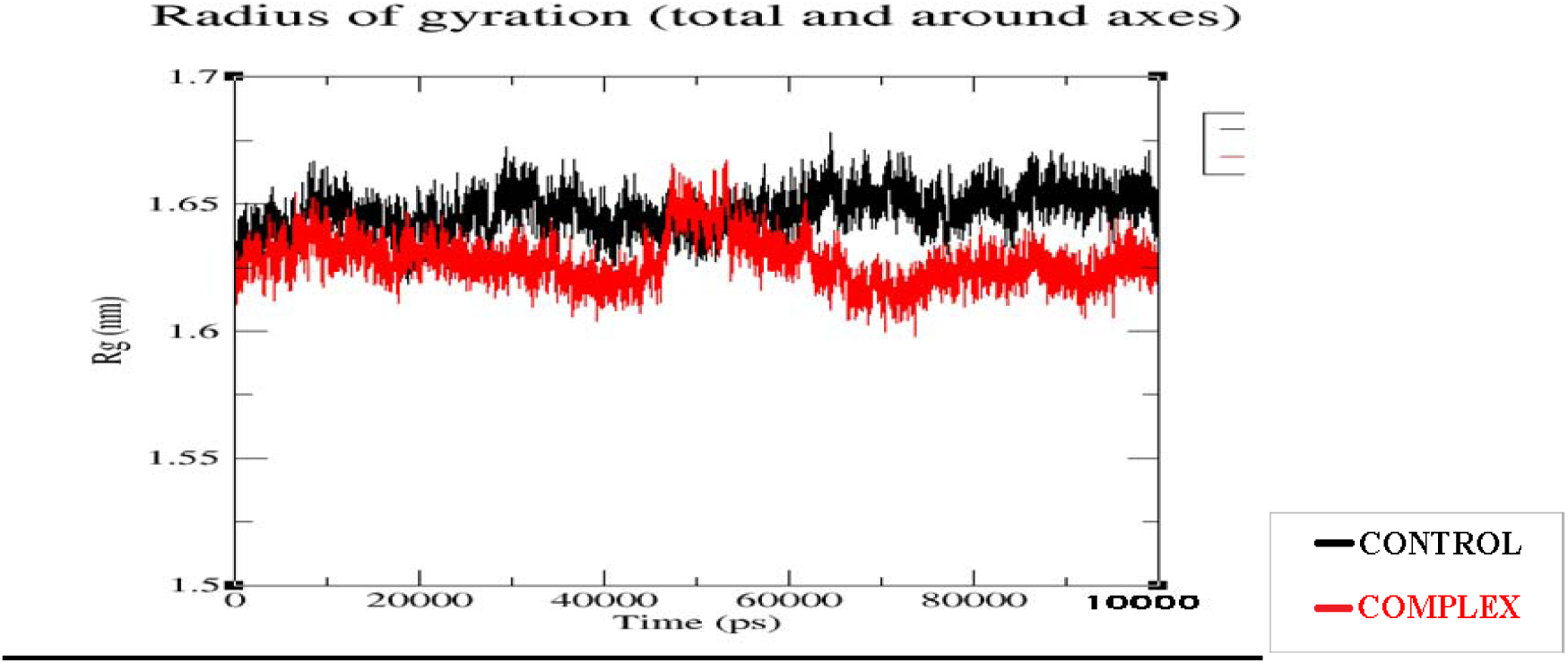
Rg plot of NDM1 – N-hydroxyfuran-2-sulfonamide complex. There is an initial increase in Rg initially in both control and complex followed by relative stability throughout the remaining duration of simulation with some minimal fluctuation. The Rg of compl x is slightly less than that of control because of the confirmational changes induced due to binding. This data suggests that the complex retains its compactness a as Rg ranges within 1.60nm to 1.67nm throughout the duration of simulation which is not a significant fluctuation.

**Fig 11.**
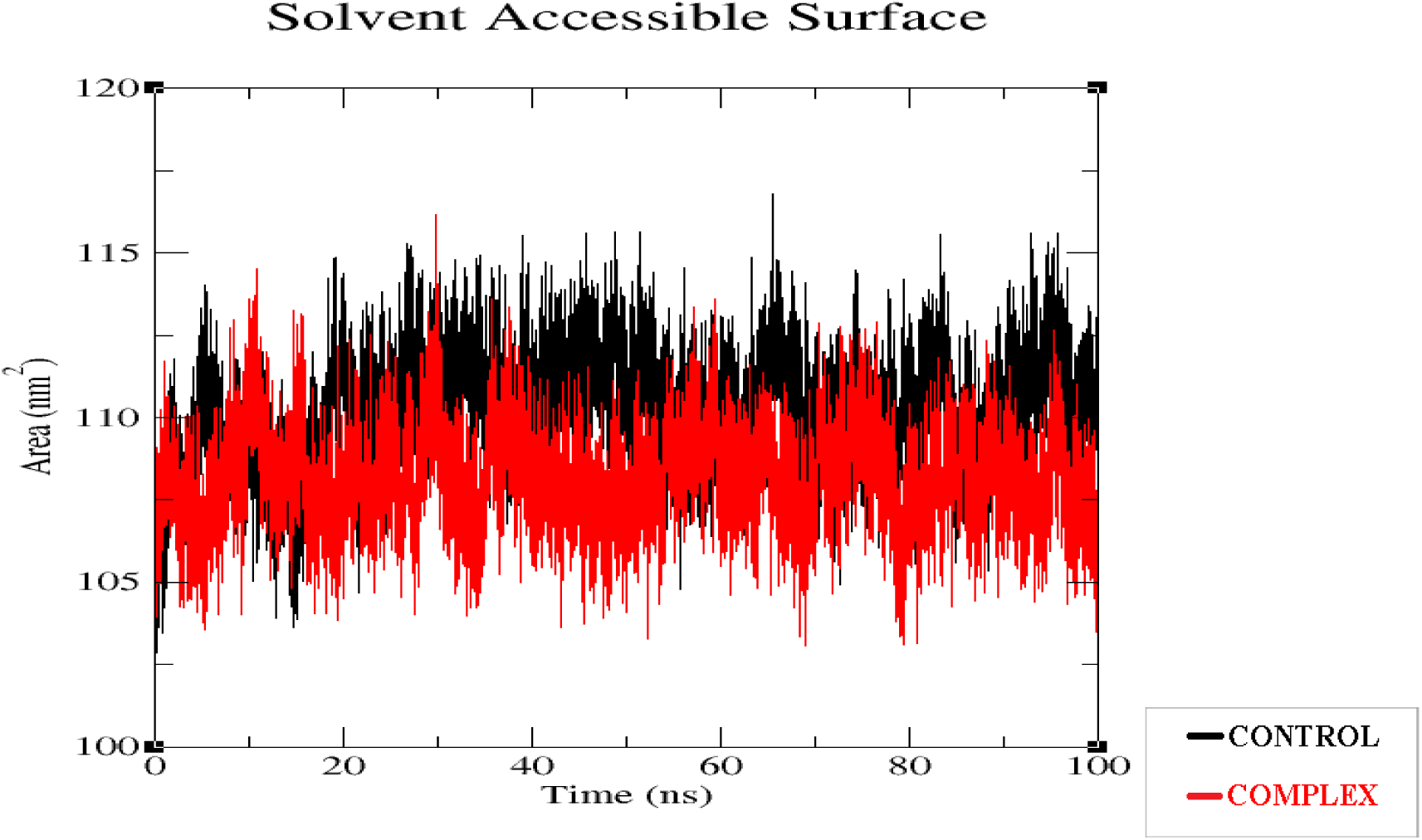
SASA plot of NDM1 – N-hydroxyfuran-2-sulfonamide complex. There is an initial fluctuation in SASA from 102nm^2^ to 112nm^2^.This might be because of ligand binding and confirmational changes. It is followed by a stable phase where SASA ranges between 105nm^2^ to 110n ^2^ in both control and the complex. SASA in complex is slightly higher than that of control due to the changes it has undergone but the fluctuation is not a significant one which leads to a conclusion that the complex does not undergo major structural rearrangements for the duration of simulation indicating the complex is stable for ligand to act on the protein.

## 4. CONCLUSION

The main aim of the research was to carry out *in silico* drug discovery and to carry out Docking studies NDM – 1 protein. The protein NDM-1 D199N with Compound 16(pdb id 6ny7) was downloaded from pdb database. The native ligand in this protein was identified as [(5,7-dibromo-2-oxo-1,2-dihydroquinolin-4-yl) methyl] phosphonic acid whose structure was downloaded from pubchem database to conduct docking studies. The obtained interactions from the docking studies was compared with the standard interactions in the pdb database and the docked results were found to be within acceptable limits. Docking of standard molecules was conducted with the same active site in the protein using PyRx and the results were analysed using Discovery studio visualizer. On analysing the results, Sulfonamide was choosen as the molecule for virtual screening. Large number of sulfonamide molecules were found in pubchem database which were filtered to about 1400 using on Verber’s and Lipinski’s rule of drug likeliness. All these molecules were docked with the active site in the protein using PyRx. The top 5% of these molecules (∼60) were selected as top hits for further study.

ADMET studies of these 60 molecules was performed using online servers ADMETLab2.0 and Pro-Tox-II followed by docking studies using Arguslab software. On analysing the obtained results, It was found that most of the molecules interacted with residues HIS:250, ASN:220, VAL:73, LYS:211.

Among these, **(3Z)-N-hydroxypenta-1,3-diene-2-sulfonamide** (Pubchem CID 118156306) and **N- hydroxyfuran-2-sulfonamide** (Pubchem CID 46175386) were found to be least toxic and had better ADME properties compared to other compounds. Molecular dynamics simulations were carried out on the complex formed these ligands with the protein and the obtained data showed that the complexes were stable for sufficient duration to act on the target protein. By the findings of this study, It was concluded that these two compounds have high chances of being developed as a drug for treating diseases and infections caused by *K.pneumoniae* producing NDM-1 strains. Further development of the compounds would be performed in future.

## ACKNOWLWGEMENT

We acknowledge and thank all the staff of Department of Pharmaceutical Chemistry, Mallige college of Pharmacy and Department of Life Sciences, Garden City University for guiding and assisting us with the research.

## Supplement file

All sulfonamide compounds screened against the target NDM-1 protein

- All other supplement files and information of the study are stored in github repository.

Available at: https://github.com/Raghu-102433/Insilico-Drug-Discovery-for-NDM-1-Beta-Lactamase

## REFERENCES

1. World Health Organization 2023, Antimicrobial Resistance, World Health Organization, World Health Organization.[online] Available at: https://www.who.int/news-room/fact-sheets/detail/antimicrobial-resistance.

2. Worthington RJ, Bunders CA, Reed CS, Melander C. Small molecule suppression of carbapenem resistance in NDM-1 producing Klebsiella pneumoniae. ACS medicinal chemistry letters. 2012 May 10;3(5):357–61.

3. Chen X, Li L, Chen S, Xu Y, Xia Q, Guo Y, Liu X, Tang Y, Zhang T, Chen Y, Yang C. Identification of inhibitors of the antibiotic-resistance target New Delhi metallo-β-lactamase 1 by both nanoelectrospray ionization mass spectrometry and ultrafiltration liquid chromatography/mass spectrometry approaches. Analytical chemistry. 2013 Aug 20;85(16):7957–65.

4. Khan AU, Maryam L, Zarrilli R. Structure, genetics and worldwide spread of New Delhi metallo-β-lactamase (NDM): a threat to public health. BMC microbiology. 2017 Dec;17:1–2.

5. Li X, Wang Q, Zheng J, Guan Y, Liu C, Han J, Liu S, Liu T, Xiao C, Wang X, Liu Y. PHT427 as an effective New Delhi metallo-β-lactamase-1 (NDM-1) inhibitor restored the susceptibility of meropenem against Enterobacteriaceae producing NDM-1. Frontiers in Microbiology. 2023 Apr 17;14:1168052.

6. Ekins S, Mestres J, Testa B. In silico pharmacology for drug discovery: applications to targets and beyond. British journal of pharmacology. 2007 Sep;152(1):21–37.

7. Fernández L, Hancock RE. Adaptive and mutational resistance: role of porins and efflux pumps in drug resistance. Clinical microbiology reviews. 2012 Oct;25(4):661

8. Nikaido H, Takatsuka Y. Mechanisms of RND multidrug efflux pumps. Biochimica et Biophysica Acta (BBA)-Proteins and Proteomics. 2009 May 1;1794(5):769–81.

9. Pemberton OA, Jaishankar P, Akhtar A, Adams JL, Shaw LN, Renslo AR, Chen Y. Heteroaryl phosphonates as noncovalent inhibitors of both serine-and metallocarbapenemases. Journal of medicinal chemistry. 2019 Sep 4;62(18):8480–96.

10. Hutchison GR, Morley C, James C, Swain C, De Winter H, Vandermeersch T, O’Boyle NM. Open Babel Documentation.

11. Morris GM, Goodsell DS, Pique ME, Lindstrom W, Huey R, Forli S, Hart WE, Halliday S, Belew R, Olson AJ. Automated Docking of Flexible Ligands to Flexible Receptors. user guide AutoDock. 2010.

12. Shaker B, Yu MS, Lee J, Lee Y, Jung C, Na D. User guide for the discovery of potential drugs via protein structure prediction and ligand docking simulation. Journal of Microbiology. 2020 Mar;58:235–44.

13. SlideShare. (2024). Protein modeling with discovery studio. [online] Available at: https://www.slideshare.net/slideshow/protein-modeling-with-discovery-studio-slides/8397277.

14. Xiong G, Wu Z, Yi J, Fu L, Yang Z, Hsieh C, Yin M, Zeng X, Wu C, Lu A, Chen X. ADMETlab 2.0: an integrated online platform for accurate and comprehensive predictions of ADMET properties. Nucleic acids research. 2021 Jul 2;49(W1):W5–14.

15. Banerjee P, Eckert AO, Schrey AK, Preissner R. ProTox-II: a webserver for the prediction of toxicity of chemicals. Nucleic acids research. 2018 Jul 2;46(W1):W257–63.

16. Justin AL. From proteins to perturbed Hamiltonians: a suite of tutorials for the GROMACS-2018 molecular simulation package [article v1. 0]. Living Journal of Computational Molecular Science. 2018;1(1):5068.

17. Abraham M, Alekseenko A, Basov V, Bergh C, Briand E, Brown A, et al. GROMACS 2024.4 Manual. Zenodo; 2024.

18. A. Bondi, vander waals Volumes and radii, J. Phys. Chem. 68(1964) pp. 441–451.

